# Blood, sweat and tears: a review of non-invasive DNA sampling

**DOI:** 10.1101/385120

**Authors:** M.C. Lefort, R.H. Cruickshank, K. Descovich, N.J. Adams, A. Barun, A. Emami-Khoyi, J. Ridden, V.R. Smith, R. Sprague, B. Waterhouse, S. Boyer

## Abstract

The use of DNA data is ubiquitous across animal sciences. DNA may be obtained from an organism for a myriad of reasons including identification and distinction between cryptic species, sex identification, comparisons of different morphocryptic genotypes or assessments of relatedness between organisms prior to a behavioural study. DNA should be obtained while minimizing the impact on the fitness, behaviour or welfare of the subject being tested, as this can bias experimental results and cause long-lasting effects on wild animals. Furthermore, minimizing impact on experimental animals is a key Refinement principle within the ‘3Rs’ framework which aims to ensure that animal welfare during experimentation is optimised. The term ‘non-invasive DNA sampling’ has been defined to indicate collection methods that do not require capture or cause disturbance to the animal, including any effects on behaviour or fitness. In practice this is not always the case, as the term ‘non-invasive’ is commonly used in the literature to describe studies where animals are restrained or subjected to aversive procedures. We reviewed the non-invasive DNA sampling literature for the past six years (380 papers published in 2013-2018) and uncovered the existence of a significant gap between the current use of this terminology (i.e. ‘non-invasive DNA sampling’) and its original definition. We show that 58% of the reviewed papers did not comply with the original definition. We discuss the main experimental and ethical issues surrounding the potential confusion or misuse of the phrase ‘non-invasive DNA sampling’ in the current literature and provide potential solutions. In addition, we introduce the terms ‘non-disruptive’ and ‘minimally disruptive’ DNA sampling, to indicate methods that eliminate or minimise impacts not on the physical integrity/structure of the animal, but on its behaviour, fitness and welfare, which in the literature reviewed corresponds to the situation for which an accurate term is clearly missing. Furthermore, we outline when these methods are appropriate to use.

## 1. INTRODUCTION

DNA data are becoming increasingly important in animal biology [1], both for experimental and observational studies. This is partially driven by the progressively cheaper and more user-friendly ways of accessing genomic information [2]. Analysis of genetic material provides data for myriad uses. In addition to analysis of phylogenetic relationships or population genetics, DNA analysis is required to determine basic information about individuals of many species [3]. When DNA analysis is required for purposes such as sexing, kinship and differentiation between cryptic species prior to experimentation with the same individuals, the DNA sampling procedure could bias the results of the subsequent experiment. It is therefore essential to minimise the effect that DNA sampling can have on the fitness or behaviour of the subject being tested. Furthermore, ethical use of animals in experimentation is guided by the ‘3Rs’ framework of Refinement, Replacement and Reduction (e.g. [4]). The impact of DNA collection is particularly relevant to the principle of Refinement where techniques with the lowest impact on the animal model should be used whenever possible. Refinement of experimentation is only possible when impact on the animal is accurately identified.

Methods of DNA collection were originally defined as ‘non-invasive’ if “*the source of the DNA is left behind by the animal and can be collected without having to catch or disturb the animal*” [5, 6], for example when genetic material was left behind in traces or scats (i.e. *sensu* environmental DNA (eDNA)), implicitly avoiding any impact on animal welfare.

These non-invasive DNA sampling procedures have been applied to study a wide range of animal taxa and answer various questions such as species identification, sexing, population genetics, description of the diet etc. To draw a comprehensive picture of the current use of these methods, we conducted a systematic review of the recent literature (2013-2018) and discuss what non-invasive DNA sampling is used for as well as issues relating to the misuse of the term.

## 2. METHOD

We conducted a keyword-based search on the Web Of Science core collection using the keywords DNA and non-invasive or DNA and noninvasive, as both spellings were originally proposed and are in common use [5, 6]. We restricted our search to articles published in relevant disciplines and between 2013 and 2018. The search command used was the following:

(TS=((dna AND non-invasive) OR (dna AND noninvasive)) AND SU=(ecology OR zoology OR ornithology OR environmental sciences OR entomology OR fisheries OR behavioural science OR Biodiversity & Conservation) AND PY=(2013 OR 2015 OR 2017 OR 2014 OR 2016 OR 2018))

Results were then refined to experimental papers written in English. On the 21st of August 2019, this search yielded 429 articles. We screened these articles retaining those in which animal DNA samples were actually collected, leading to 397 articles, and removed articles with insufficient methodological information to draw conclusions about the specific questions investigated. A total of 380 papers were retained in our final dataset (see list in Supplementary Table 1). Although this dataset may not be exhaustive; it is taken to be representative of the current literature on non-invasive DNA sampling.

During the same time period and in the same fields as above, we estimated the total number of articles focusing on invertebrates versus vertebrates using the following commands:

- (TS=(mammal) OR TS=(vertebrate) OR TS=(bird) OR TS=(amphibian) OR TS=(reptile) OR TS=(fish) NOT (TS=(insect) OR TS=(invertebrate) OR TS=(crustacean) OR TS=(annelid) OR TS=(echinoderm) OR TS=(nemathelminth) OR TS=(arachnid) OR TS=(arthropod) OR TS=(plathelminth)) AND SU=(ecology OR zoology OR ornithology OR ecology OR environmental sciences OR entomology OR fisheries OR behavioural science OR Biodiversity & Conservation) AND PY=(2013 OR 2015 OR 2017 OR 2014 OR 2016 OR 2018))
- (TS=(insect) OR TS=(invertebrate) OR TS=(crustaceans) OR TS=(annelid) OR TS=(echinoderm) OR TS=(nemathelminth) OR TS=(arachnids) OR TS=(arthropod) OR TS=(plathelminth) NOT (TS=(mammal) OR TS=(vertebrate) OR TS=(bird) OR TS=(amphibian) OR TS=(reptile) OR TS=(fish)) AND SU=(ecology OR zoology OR ornithology OR ecology OR environmental sciences OR entomology OR fisheries OR behavioural science OR Biodiversity & Conservation) AND PY=(2013 OR 2015 OR 2017 OR 2014 OR 2016 OR 2018))

The results from these searches were used as non-exhaustive but comparable numeric estimates only, and were therefore not further curated. The abstract and the method section of each papers were carefully screened to check whether the methods used complied with the original definition proposed by Taberlet et al.[6] or not. A middle-ground category, labelled as “potentially affecting territory”, was created for cases where faecal samples were taken from wild animals that are known to use dejections as territory or social marking. We excluded from this category, studies that specifically mentioned only partial collection of faeces. Where multiple methods were used in the same study, these were classified as compliant with the definition by Taberlet et al. only if all the methods used were compliant or if invasive sampling methods were clearly identified from non-invasive ones. The latter required screening of the whole paper.

Statistical analyses were conducted with R[7] (version 3.6) and RStudio[8] (version 1.2.1335). Packages used included *stats*, *googleVis* and *bipartite*. Statistical significance was set at 5%.

## 3. WHAT NON-INVASIVE DNA SAMPLING IS USED FOR

Our systematic review captured 380 articles for which samples were collected from 96 different countries on all continents except mainland Antarctica (Fig 1a). The number of papers detected per year was stable between 2013 and 2018 (X^2^ = 4.421, df = 5, p-value = 0.4877). The sampling methods used varied between 2013 and 2018 (X^2^ = 39.754, df = 25, p-value = 0.03091), with in particular an increase in the use of eDNA (Fig 1b).

**Figure 1.**
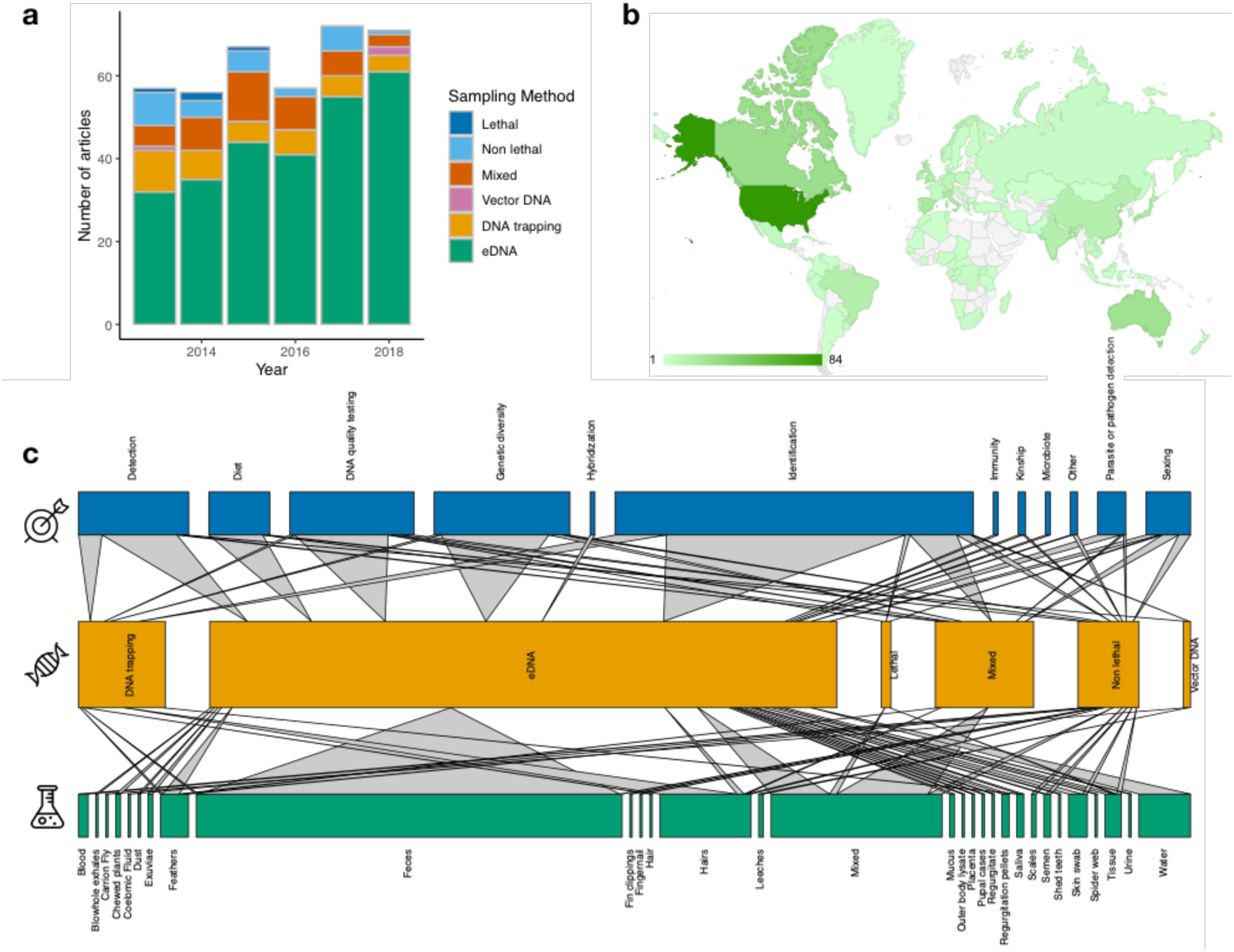
Summary statistics of the literature review on the use of “non-invasive DNA sampling” between January 2013 and December 2018 (n=380). **a:** Number of articles in relation to the sampling method used between 2013 and 2018. **b**: Countries of origin of the samples analysed in the reviewed papers. Countries in grey were not represented in our review, countries coloured in various shades of green provided samples for 1 to 84 of the reviewed papers (see in-graph legend for colour scale). **c**: Bipartite network of the main aim of the studies in blue, the type of sampling method used in orange (see Table 1 for definitions) and the nature of the samples collected in green. The horizontal width of the rectangles is proportional to the number of articles in each category.

Among the studies captured in our review, 40% aimed at identifying organisms at the species level, for example to produce biodiversity inventories, or at the individual level (Fig 1c). The latter was often conducted in the context of Capture Mark Recapture (CMR) studies (e.g.[9]), where it is essential to identify individuals. Individual genotyping was also often attempted to measure genetic diversity or for population genetic studies (e.g.[10]) in 15% of the reviewed articles. The development of new protocols where the quality of the DNA obtained non-invasively was the center of interest was the aim in another 14% of the studies. Other recurrent foci were on the detection of presence (12%), the study of animals’ diet (7%) or the sexing of individuals (5%).

The type of samples collected varied widely and 30 different categories were recorded. However, a large number of the studies focused on faecal samples collected as eDNA (48%) (Fig 1c). Another 19% of studies were based on the collection of more than one type of samples, often including faeces. Hair samples, water samples and feathers were the next most represented sample types in our dataset (10%, 6% and 3% of studies respectively). Hair samples were mainly collected through DNA trapping, while feather and water samples were generally collected using an eDNA approach. We also uncovered a variety of much more atypical sample types such as insect pupal cases, urine, fingernails, placenta, mucus etc.

Overall, the substantial majority of sampling methods (71%), were based on the collection of eDNA, while DNA trapping was rarely used (10%). Other cases included studies using several different methods (11%) and few very specific cases (Fig 1c). For example, invertebrates such as leeches^10^ and carrion flies^9^ were used to sample the DNA of the species on which they feed (Fig 1c). More surprising, a number of studies only used non-lethal (but invasive) or even lethal sampling methods (8% of the reviewed papers). Such methods are in breach of the definition of non-invasive DNA sampling as proposed by Taberlet et al.[6]. In fact, 58% of reviewed papers using the phrase “non-invasive” or “noninvasive” did not comply with this definition (Fig 2a) even when this phrase was present in the title of the article (59% of non-complying articles).

**Figure 2.**
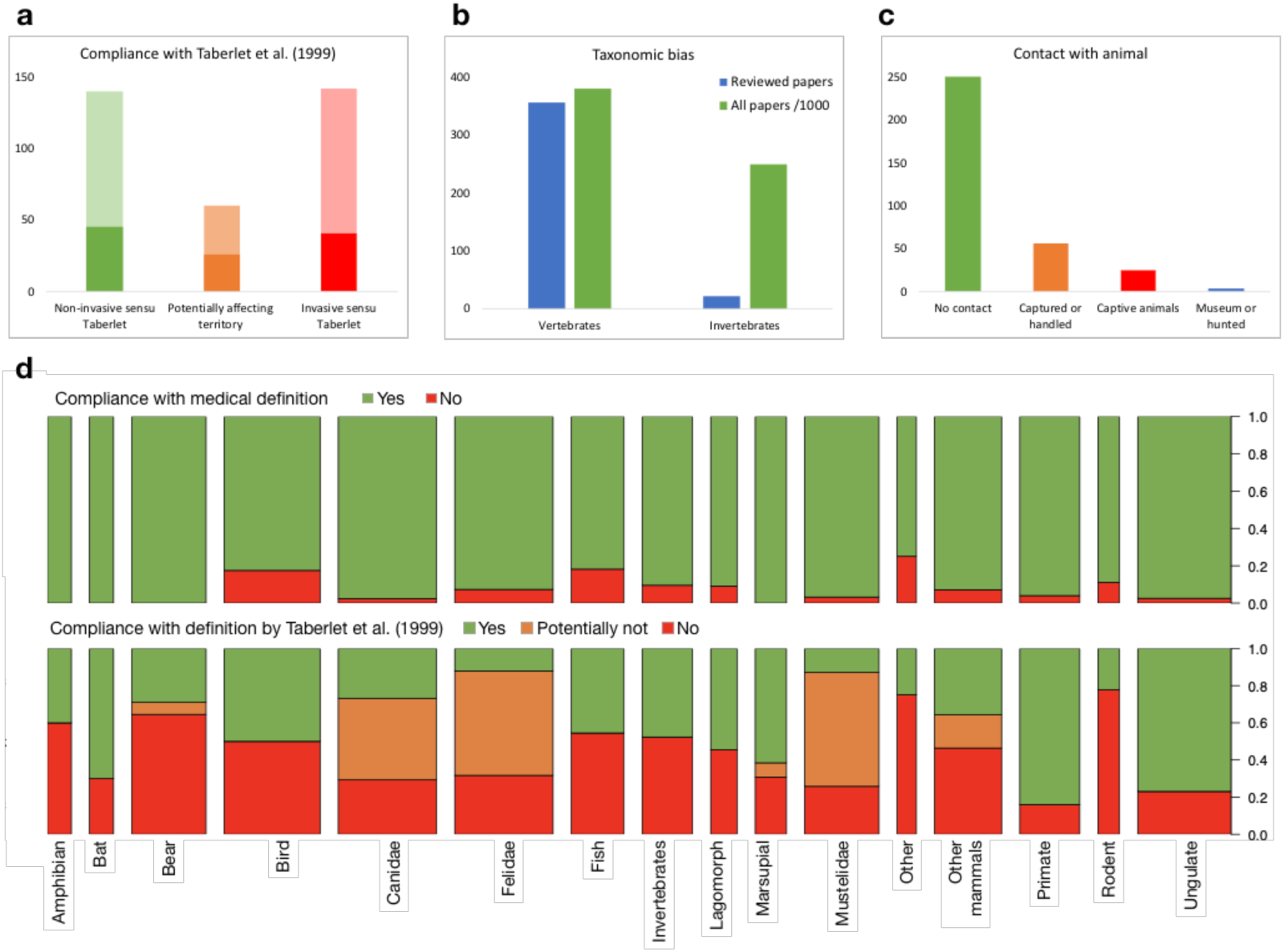
Summary statistics of the main issues exposed by our literature review on the use of “non-invasive DNA sampling” between January 2013 and December 2018 (n=380). For a, b, and c, the y-axis is the number of papers. For d, the y-axis is the proportion of papers and the width of the bars is proportional to the number of papers for each taxonomic group. **a:** Compliance of papers with the original definition proposed by Taberlet et al. ([6]). Studies where multiple methods were used (n=31) were classified as compliant with the definition by Taberlet et al. only if all the methods used were compliant OR if invasive sampling methods were clearly identified by the authors. Dark colours correspond to papers where the phrase “non-invasive” was present in the title, lighter colours correspond to papers where the phrase “non-invasive” was not present in the title. The orange bar (labelled as “potentially affecting territory”, corresponds to cases where territory marking and social interactions may have been affected by the removal of faecal samples. **b**: Taxonomic bias in the non-invasive DNA sampling literature. Number of papers reviewed that focus on invertebrates or vertebrates compared to all papers on invertebrate or vertebrate (see Method section for search command). **c**: Number of papers complying (in green) or not complying with the no contact criteria proposed by Taberlet et al. ([6]), because animals were either captured or handled for DNA sampling (orange), held in captivity (red) or had been killed (blue). **d**: Proportion of papers complying with different definitions of non-invasive sampling in relation to the taxonomic group studied. Top: compliance with the common definition of a non-invasive medical or veterinary procedure, (i.e. one not involving puncture of the skin or other entry into the body ([70]). Bottom: compliance with the definition of non-invasive DNA sampling proposed by Taberlet et al. ([6]). Orange boxes (labelled as “Potentially Not”) correspond to cases where territory marking and social interactions may have been affected by the removal of faecal samples.

## 4. DNA COLLECTION AND THE NON-INVASIVE MISNOMMER

Subsequent to its original definition, the term non-invasive has often been misapplied in the literature [11]. In practice, so-called ‘non-invasive’ methods have often encompassed DNA collection techniques that preserve the physical integrity of an organism but have an unmeasured, and potentially significant, impact on the fitness, behaviour or welfare of the subject being studied. For example, the following DNA collection methods were all defined as ‘non-invasive’ by the respective authors: gentle pressure applied to the thorax and abdomen of carabid beetles (*Poecilus cupreus*) to trigger regurgitation [12]; flushing of sage-grouse (*Centrocercus urophasianus)* from their roost sites to collect fresh faecal pellets [13]; and trapping, handling and cloacal swabbing of lizards (*Phrynosoma cornutum*) [14]. Misleading use of terminology in biology and ecology is a longstanding concern [15–17]. To demonstrate the extent of the issue, we conducted a systematic review of the recent literature (2013-2018) and evaluated how well papers using the term “non-invasive DNA sampling” complied with the original definition by Taberlet et al. [6].

When the terminology for DNA sampling is misapplied as being non-invasive when it is not, readers unfamiliar with the scientific literature on DNA sampling (e.g. decision makers, conservation managers, and other end-users), may be misled in thinking that the described method can be applied without affecting the fitness nor behaviour of the target animals. Misnaming DNA sampling is also problematic for assessing impact on animals, identifying opportunities for refinement, and for judging the validity and quality of the data collected. Using more precise terminology could also help scientists realise that they may have been using invasive methods after all, and encourage them to consider reducing the impact of their sampling and/or search for truly non-invasive alternatives. The main issues exposed by our literature search are summarised in Box 1.

### BOX 1: THE SEVEN SINS OF NON-INVASIVE DNA SAMPLING

#### Sin 1: Taxonomic bias

One conspicuous result from our review was that only 18 studies (∼6% of the reviewed papers) focused on invertebrates compared to 356 focusing on vertebrates (Fig 2b). This striking imbalance implies that non-invasive methods are rarely considered for sampling invertebrate DNA. When authors claimed to use non-invasive DNA sampling on invertebrates, they failed to do so in 55% of the cases (Fig 2d), and even used methods that alter the physical integrity of the organism in 10% of the cases. For example, Rorat et al. [18] collected individual earthworms, which they then electrified “*lightly*” to induce coelomic secretion. Yet, truly non-invasive methods exist for invertebrates, for example through field collection of insect exuviae [19], pupal cases [20], empty mummies [21], dust [22], soil[23], or water samples[24].The misuse of the term non-invasive DNA sampling also varies in relation to the taxonomic group of interest within vertebrates (Fig 2d) (X^2^ = 190.69, df = 30, p < 2.2e-16). For example, 27% of the studies on fish involved alteration of the physical integrity of the organism. These included fin clipping in eels (*Anguilla anguilla*) [25] and sting amputation in rays (*Aetobatus narinari*) [26] which were both considered non-invasive because these body parts can regenerate, despite the fact that fin clipping is known to be painful for fish [27].In comparison, less than 4% of the studies focusing on mammals, involved biopsies.

#### Sin 2: Misclassification of faeces as non-invasive DNA samples

The majority of the literature on non-invasive DNA sampling included the collection of faecal samples (62% of all studies reviewed here). Faecal collection is very prevalent in the field and assumed to be non-invasive by most authors. However, our analysis shows that 47% of the studies focusing solely on faecal sampling did not comply with the original definition of non-invasive DNA sampling. This included detection of animals and collection of faecal samples using aircraft (e.g. [28]), which may increase stress in animals (e.g. [29]) or cases where animals were being held in captivity (e.g. [30]), specifically captured to obtain faecal samples (e.g.[31]). For example, Jedlicka et al. [32] “extracted DNA from noninvasive fecal samples” of Western Bluebirds (*Sialia mexicana*) by catching adults and placing them in brown paper bags. Despite focusing on faecal samples, these procedures do not fit the definition proposed by Taberlet et al. [6]. The central misconception, here is that there is no such thing as “non-invasive DNA samples”. Rather than the type of sample, it is the method of sampling that needs to be scrutinized for its invasiveness. Another key issue with faecal sampling is that many animals mark their territory using faeces to dissuade potential intruders (e.g. in wolf communities, see [33]) and also use such marks to recognise individuals from neighbouring territories, avoid unnecessary conflict and promote non-agonistic social encounters such as mating. Therefore, even when collected opportunistically after the animal has left, faecal sampling can in some cases affect the marking behaviour of territorial species (e.g. [34]) (Fig 2a).

#### Sin 3: Baiting DNA traps

In most studies using a DNA trapping strategy (90%), researchers employed bait or lures to increase the yield of their traps. Very few studies used non-lured DNA traps, for example, barb wire placed at sites used by brown bears (*Ursus arctos*) [43, 44] or modified body snares at otter (*Lontra canadensislatin*) latrine sites, to collect hair [45]. Although it seems perfectly legitimate (and often essential) to increase the attractiveness of DNA traps with food [46], scent marks from other individuals [47] or other attractants (e.g. Valerian essence for cats) [48], the animal’s behaviour will obviously be modified as a consequence and therefore, these methods cannot be considered fully non-invasive sensu Taberlet et al. [6].

#### Sin 4: Combining invasive and non-invasive methods

In a few examples the impact of the sampling strategy on animal behaviour is obvious from the article’s title itself, for example when baited traps are mentioned (e.g. [48]). However, in many more papers (n=35) confusion arises because authors used the phrase “non-invasive sampling” or “non-invasive DNA sampling” while a variety of sampling techniques were actually applied, some of which were non-invasive and some of which were invasive sensu Taberlet et al. [6]. This lack of clarity about what is non-invasive and what is not can be misleading for the reader. Some authors clearly stated the invasiveness of the different methods used (e.g. [49–51]), however, most papers where mixed DNA sampling strategies were applied did not specify which of these methods were considered non-invasive.

Another facet of this issue arises when tools (e.g. new primers, extraction protocols, DNA conservation methods) are developed specifically for analysing samples collected non-invasively but are actually tested only (or partly) on samples that were collected invasively (n=17) for example by capturing animals to perform the sampling (e.g. [52, 53]). It is essential in such cases that authors fully acknowledge the invasiveness of the sampling method(s) they used. Often this is not clearly specified.

#### Sin 5: A bird in the hand is no better than two in the bush

Trapping and restraint of wild animals is recognised as a significant stressor that can result in distress, injury, and death (e.g. [54]). Capturing and/or handling animals for DNA sampling was observed in 24% of all articles reviewed here (Fig 2c), despite the clear definition given by Taberlet et al. [6] that non-invasive DNA is “*collected without having to catch or disturb the animal*”. Indeed, capture and/or handling of individuals to obtain DNA samples (e.g. saliva swabbing) can induce long-lasting stress effects [55, 56], and there are very few cases where capturing an animal might have no effects on its future behaviour. Therefore, when animals must be held captive, transported or restrained in order to perform DNA sampling, the method cannot meet the definition of non-invasive DNA sampling *sensu stricto* [6]. Skin swabbing of octopus (*Enteroctopus dofleini*) for example [57], is unlikely to be possible in the wild without disturbing the animal and the potential negative impacts on animal welfare (see [58] for a review on cephalopod welfare) must still be recognised.

Another common scenario where the animals are held during DNA sampling relates to the use of museum specimens or animals that were killed for other purposes (n=4). Whether they were legally hunted or poached and confiscated (e.g.[59]), this type of sampling does not qualify as non-invasive due to the disturbance and/or death of the animal through human activity. Often, a better term for such sampling is “non-destructive”, which does not damage the specimen [60, 61] (Table 1). On the other hand, tissue sampling from animals that were found dead of natural causes is analogous to eDNA left behind by a free ranging animal and can be considered non-invasive (e.g. [62]). It should be noted, however, that opportunistic sampling from animals already killed for other purposes (e.g. culling, museum samples) may be an ethical option because it reduces the need to otherwise target living animals and conforms to the principle of Reduction (reducing the number of affected animals) under the 3Rs framework.

**Table 1.** Glossary of terms as used in this review.

| Term | Definition |
| --- | --- |
| **DNA trapping** | Remotely obtaining DNA from one or more unknown individual organisms by taking a sample while they are present. This usually involves some sort of trap or device, which may or may not be disruptive. |
| **eDNA sampling** | Obtaining trace DNA left behind by one or more unknown organisms, by sampling the environment when those organisms are no longer present at the point of sampling. |
| **Minimally disruptive DNA sampling** | Obtaining DNA with minimised effects on the animal's fitness, behaviour and welfare. To a minimised extent, such method may affect the structural/physical integrity of the organism. |
| **Minimally invasive DNA sampling** | Obtaining DNA with minimised effects on the animal's structural/physical integrity. To a minimised extent, such method may affect the behaviour and welfare of the organism. |
| **Non-destructive DNA sampling** | Obtaining DNA from a known individual organism in such a way that the organism may be killed, but not destroyed, so that it can be preserved as a voucher specimen. |
| **Non-disruptive DNA sampling** | Obtaining DNA without affecting the animal's fitness, behaviour and welfare. |
| **Non-invasive DNA sampling *sensu lato*** | Obtaining DNA without affecting the physical integrity of the animal's through puncturing the skin or other entry into the body (derived from the medical definition of a non-invasive procedure). |
| **Non-invasive DNA sampling *sensu stricto*** | Obtaining DNA that was left behind by the animal and can be collected without having to catch or disturb the animal (from Taberlet et al. 1999) |
| **Non-invasive procedure** | A procedure that does not involve puncture of the skin or other entry into the body (such as use of an endoscopic device). |
| **Non-lethal DNA sampling** | Obtaining DNA from an organism in such a way that the organism is not killed. This broad category includes invasive and non-invasive methods. |

#### Sin 6: All or nothing

Only 42% of the reviewed studies fully met the criteria of the original definition of non-invasive DNA sampling. In most cases, however, authors tried to minimise the impact of sampling, but the nature of the definition proposed by Taberlet et al. [6] leaves no middle ground between invasive and non-invasive sampling methods. One potential solution to this is to use the term “minimally-invasive DNA sampling”, which can be defined as obtaining DNA with minimised effects on the animal’s structural/physical integrity, and potential impact on the behaviour and welfare of the organism (Table 1). In our dataset, this term was used in six studies to qualify skin swabbing of fish [63], amphibians [64] and bats [65], feather plucking of gulls [66], cloacal swabbing in rattlesnakes [67] and ear biopsies in rodents [68]. A broader use of this term would lead to more accurate reporting, for which potential impacts of the sampling are acknowledged, while still emphasising the aspiration of the authors to minimise those impacts. The challenge associated with the use of such a term would be to define where ambiguities fall between minimally-invasive and invasive sampling methods.

#### Sin 7: Equating a non-invasive procedure with non-invasive DNA sampling

The lack of perceived stress or pain experienced by an animal is often used as a criterion to support the classification of a method as non-invasive. For example, du Toit et al. [69] stated that “*Pangolin scales consist of non-living keratin, therefore taking scale clippings is considered to be non-invasive*”. This statement relates to the common definition of a “non-invasive” medical or veterinary procedure, i.e. one that does not involve puncture of the skin or other entry into the body [70]. This definition (rather than the one by Taberlet et al. [6]) seems to be the one adopted by most authors (93% of the reviewed papers complying) (Fig 2d). This was also the case for several articles at the frontier between medical/veterinary fields. Kauffman et al. [71] for example, called the sampling of vaginal swabs and urine from captive dogs non-invasive. Similarly, Reinardy et al. [72] designated as ‘non-invasive’ a procedure consisting of “*lightly anaesthetizing fish and applying a slight pressure on their abdomen to expel sperm*”, which was then used for DNA analysis. These examples were rare in our dataset (n=3) probably because of our strict selection of articles from non-medical and non-veterinary domains (see selected fields in section 2). Nonetheless, as science becomes increasingly transdisciplinary and genetic methods developed in neighbouring fields are used in ecology, this type of confusion is likely to become more prevalent in the future. The discrepancy with the common definition of a non-invasive procedure comprises a significant limitation of the phrase non-invasive DNA sampling as defined by Taberlet et al. [6], and importantly, could minimise the perceived impacts of sampling methods on animal welfare, even if these impacts are significant in reality. Although this issue was first highlighted in 2006 by Garshelis who stated that: “*the term noninvasive has 2 distinct meanings, 1 biological and 1 generic, which have become intertwined in the wildlife literature*” [11], the confusion continues to riddle the current literature.

## 5. INTRODUCING THE TERMS NON-DISRUPTIVE AND MINIMALLY DISRUPTIVE DNA SAMPLING

In order to clarify some of the existing discrepancies exposed by our literature review, we propose the introduction of the term, ‘non-disruptive DNA sampling’. This term emphasises the effects of the sampling method not on the physical integrity/structure, but on the fitness and behaviour of the organism from which the sample is obtained. We define ‘non-disruptive DNA sampling’ as obtaining DNA from an organism without affecting its fitness, or causing any behaviour or welfare impact that may last longer than the duration of the sampling (Table 1). We define ‘minimally disruptive DNA sampling’ as any sampling method that minimises impacts on fitness, behaviour and welfare. Non-disruptive DNA sampling can be differentiated from ‘non-invasive DNA sampling’ which in the current literature, largely focuses on whether the method of sampling impacts physical structures of the animal (Fig 2d). The introduction of ‘non-disruptive DNA sampling’ terminology provides a functional term that appropriately focuses on the impact to the individual and not on a specific quality of the methodology (e.g. whether a physical structure is altered). We acknowledge that very few current DNA sampling methods may be entirely non-disruptive, and recommend that researchers aim at minimising disruption through protocol refinement. This could be achieved by testing the potential effects of different DNA sampling methods on survival, stress, behaviour and reproductive success as proxies for fitness. In order to make our intended meaning clear, we overlaid existing DNA sampling terms in relation to non-disruptive DNA sampling methods in the following paragraphs and in Figure 3. Rather than debating and refining existing terms, the essential point of Figure 3 is to distinguish between disruptive methods, which are likely to cause lasting effects on the behaviour, welfare or fitness of an organism, and non-disruptive ones, which do not.

**Figure 3.**
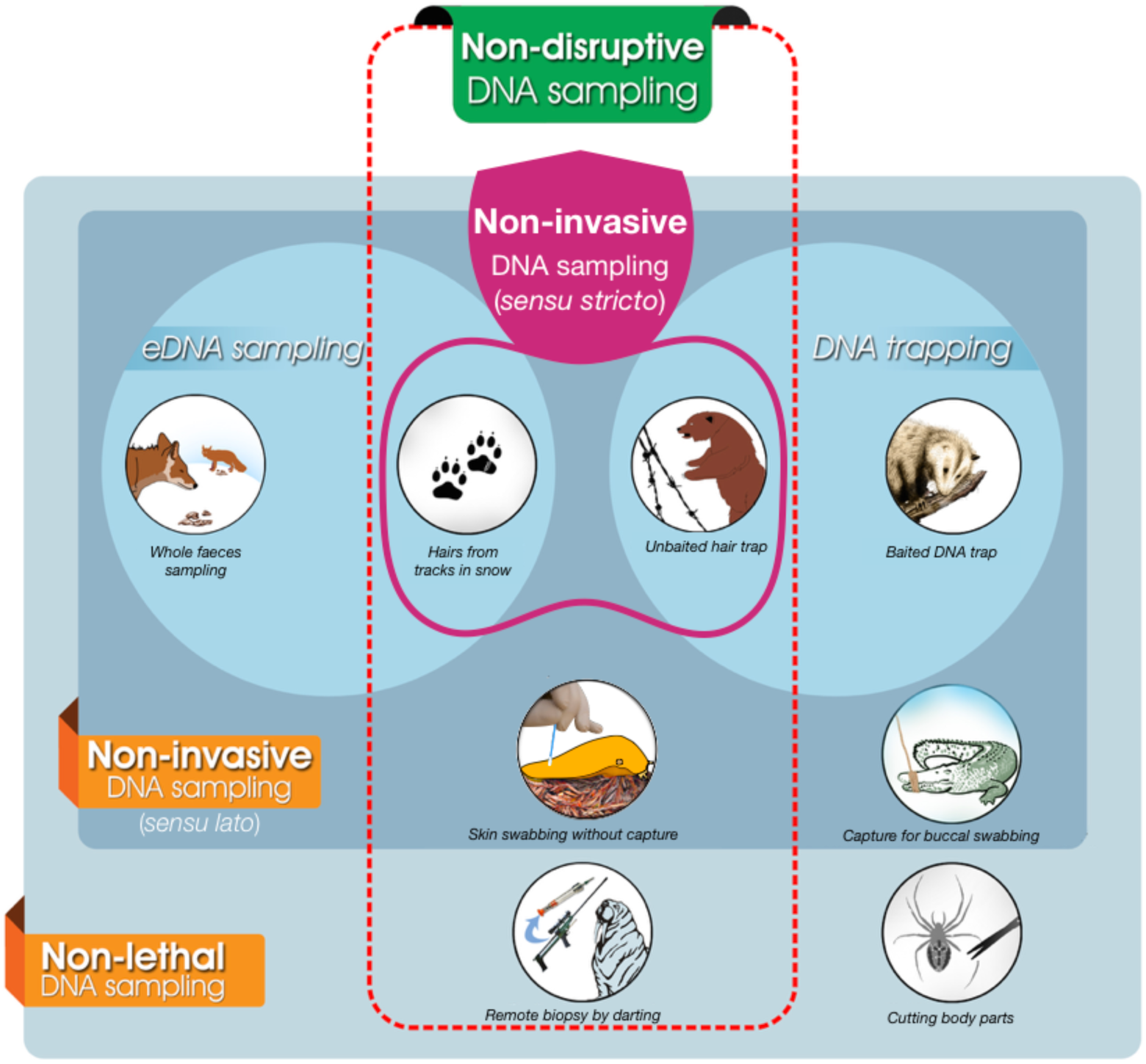
The relationship between non-disruptive, non-invasive and non-lethal DNA sampling methods. Non-invasive DNA sampling *sensu stricto* corresponds to the definition given by Taberlet et al. ([6]), Non-invasive DNA sampling *sensu lato* corresponds to the medical definition ([70]). Pictograms represent a non-exhaustive list of examples for which references are given below. From left to right and top to bottom: whole faeces sampling for species that use faecal territory marking ([113]), hairs collected in snow ([50]), hairs collected with unbaited barbed wire([43]), DNA trap baited to attract animals ([114]), skin swabbing in the field without capture ([94]), capture of reptiles for buccal swabbing ([115]), gun darting of big mammals to collect tissue sample([116]), biopsy on handled invertebrate ([117]).

### 5.1. Impact of DNA sampling on behaviour, fitness and welfare

Studies examining the effect of DNA sampling on behaviour, fitness and welfare are rare and their results are not always predictable. For example, the fitness consequences of DNA sampling methods, often measured using individual survival as a proxy for fitness (e.g. [73–75]), depends on the taxa sampled. Responses may vary strongly between species [76] and even between males and females of the same species. For instance, Vila *et al.* [77] showed that the non-lethal but invasive DNA sampling through leg or hind wing clipping had an effect on survivorship and reproductive behaviour of adult males of the protected moth *Graellsia isabelae,* while mid leg clipping had a negative impact on female mating success.

In particular cases, procedures to obtain DNA samples can also increase the fitness of animals. For example, supplementary feeding can have a direct positive impact on the fitness of birds [78], and this may occur when animals are attracted to DNA traps baited with food or feeding cages where animals are caught for DNA sampling (e.g. [79]). In mammals, remote DNA sampling using biopsy darts is known to cause little reaction from marine mammals when conducted correctly and is unlikely to produce long-term deleterious effects [80]. Gemmell and Majluf [81] found that in most cases New Zealand fur seals (*Arctocephalus forsteri*) recoiled from the impact and searched briefly for the assailant, but never abandoned their territory following the darting. Another study found that bottlenose dolphins (*Tursiops* spp) reacted similarly to the darting process regardless of being hit or not, suggesting that the reaction is mainly caused by ‘unexpected disturbance’ rather than biopsy [82]. No sign of long term altered-behaviours was observed, including probability of recapture. Despite this, all biopsy sampling involves some level of risk [80], and different individuals from the same species may react differently to similar stressful situations depending on gender [83] or individual physiological and psychological factors [84, 85]. With regards to animal welfare, Paris et al. [86] assessed the impact of different DNA sampling methods on individual welfare in frogs. They concluded that capture and toe clipping was significantly worse than capture and buccal swabbing in terms of the level of suffering experienced by an animal, and the detrimental impacts on survival. These examples illustrate that the level of disruptiveness of DNA sampling methods should be made cautiously and studies assessing their impact on fitness, behaviour and welfare should be encouraged prior to their use.

### 5.2. Examples of non-disruptive or minimally disruptive DNA sampling

Non-disruptive DNA sampling comprises all non-invasive DNA sampling *sensu stricto* i.e when the DNA is collected without the subjects being aware of the researcher’s presence or experiencing any detrimental effects (as suggested in Taberlet & Luikart [5]). For example, most eDNA sampling and DNA trapping methods do not require researcher and subject to be present at the same time and place. An important point of difference between these two methods is that eDNA is often collected somewhat opportunistically, while DNA trapping allows for strategic spatial distribution of sampling.

Examples of DNA trapping that are non-disruptive include remote plucking or hair trapping by means of unbaited hair snag traps [87, 88] or tape [89, 90] placed at well-used runs. Environmental DNA sampling includes field collection of faeces (e.g. [36]) as long as these do not affect territory marking (see section 3.2), DNA collection from footprints in the snow, such as those from the Swedish Arctic fox (*Vulpes lagopus*) [91], and from saliva on twigs, such as from ungulate browsing [92]. When DNA is collected in the presence of the animal, the effects of sampling can be minimised by avoiding or drastically limiting handling. For example, the swabbing of animals directly in the field with little [93] or no handling [94].

Sampling methods that are non-disruptive have many benefits for conservation science, because they are unlikely to introduce bias or experimental effect or impact on animal welfare. However, they may be limited in their applicability. The main limitations associated with eDNA and DNA trapping include low DNA quantity and quality [95], as well as potential contamination from non-target species [96]. Another limitation of DNA trapping might be the mixture of DNA from several different target individuals. In such instances, next-generation sequencing (NGS) or other post-PCR analysis (e.g. cloning, single stranded conformation polymorphism, high resolution melting, denaturing gradient gel electrophoresis) might be required to differentiate and identify the DNA of each individual.

A shift in focus from sampling methods that aim at avoiding breaches to physical structures of an organism, to non-disruptive or minimally disruptive methods, (avoiding impact on behaviour, fitness or welfare), means in some cases the most appropriate method may be invasive but results in a lower impact on the animal. For example, invertebrate antenna clipping in the natural environment breaches a physical structure but may result in no effects on survival (e.g. [75]) and may have lower impacts than collecting and removing specimen to captivity for faecal sampling or forced regurgitation.

Similarly, remote dart biopsy or flipper notching of marine mammals are often a preferred choice over stressful captures for DNA sampling because they only cause short term effect (if any) on the behaviour of the animal [97, 98]. Under our definitions, hair collection from the environment, unbaited DNA traps, skin swabbing in the field or remote darting on wild sea mammals could be considered non- or minimally disruptive (Fig 3).

## 6. WHEN IS NON-DISRUPTIVE DNA REQUIRED OR PREFERRED?

The selection of a DNA sampling method is usually a compromise between minimising welfare and ethical costs, and obtaining a quality DNA sample. DNA sampling methods where the specimen is in hand generally results in fresher and better-quality DNA, despite the potentially higher impact on animal behaviour or welfare. While the welfare of all experimental animals should be considered, when the subject is endangered or afforded legal protections there may be additional welfare and/or ethical issues surrounding the use of invasive DNA sampling techniques [73, 99]. Additionally, the test subject may be required to be alive for further testing or return to their natural habitat. If further tests involve capturing an animal for a laboratory experiment [100] or for translocation [101], then the effects of capturing and holding the organisms for DNA sampling are of less concern as individuals will need to be captured for these experiments anyway. However, stressful events can have a cumulative effect [102], therefore the potential for further exacerbation of stress by DNA sampling should be carefully considered.

The importance of considering non-disruptive DNA sampling also depends on the type of study undertaken. Below we describe experimental studies, field behavioural studies, and capture mark recapture (CMR) research, as three types of situations in which collection and use of non-disruptive DNA samples may be essential.

### 6.1. Laboratory-based experimentation

Non-disruptive DNA sampling is necessary for species identification, sexing or genotyping of individuals prior to laboratory-based experimentation where fitness and/or behavioural traits are to be assessed. For example, many species of birds are monomorphic, and can only be sexed using molecular analysis [103]. Similarly, many cryptic species complexes can only be elucidated genetically [104]. Laboratory-based behavioural or fitness studies involving cryptic or monomorphic species may therefore require DNA sexing or species identification of individuals before conducting research on them [100, 105] to ensure a balance of sex or species across different treatments. Even when species identification is not an issue, the organisms being studied may comprise different morphocryptic genotypes [105] that must be determined prior to experimentation in a way that does not affect their fitness or behaviour. One classical way to alleviate the effects of sampling on behaviour (for example when animals are collected in the wild and brought to the lab), is to allow for a recovery and acclimation period.

### 6.2. Behavioural studies in the field

The second major use of non-disruptive DNA sampling is when relatedness between individual subjects must be determined prior to a behavioural study conducted in the field. For example, social interactions in mammals are often linked to kinship and can be mediated by the physiological state of individuals [106]. The capture and handling of animals can modify their physiology [107], thereby affecting their social behaviour. Recent studies also suggest that although behaviours observed shortly after release may appear ‘normal’, stress levels may still be high and impact activity budgets [108]. Such effects may remain undetected but have significant implications for subsequent data reliability and validity.

### 6.3. Capture Mark Recapture

The effects of DNA sampling on animal behaviour may also affect the results of studies that are not directly examining behaviour or fitness. The third case when non-disruptive DNA sampling is recommended is when doing Capture Mark Recapture (CMR) studies. CMR studies using DNA tagging are often conducted to estimate population size (e.g. [109]), with the additional benefit of enabling population genetic analysis on the samples collected. Invasive or disruptive DNA sampling techniques may affect the survival rate of marked individuals, or introduce avoidance behaviours, which may cause trap avoidance, and the population size to be overestimated. For example, toe clipping combined with CMR is commonly used to estimate population abundance of amphibians [110], but toe clipping has been shown to decrease chances of frog recapture by 4 to 11 % for each toe removed [73]. Similarly, sampling methods that may increase the fitness of animals (e.g. feeding cages or baited DNA traps) could lead to previously sampled animals being more attracted than naïve ones [11, 111], thereby biasing the CMR results towards underestimating population size.

Such biases can be limited by the use of non-disruptive DNA sampling methods. Although eDNA has been used in CMR studies and is in most cases non-disruptive, it can have some limitations. The presence of mixed DNA samples and the lower quality of the collected DNA can lead to false positives where animals not captured previously are believed to be recaptured due to their DNA profile being indistinguishable from that of captured animals [112]. Because of this, non-disruptive DNA sampling may provide an appropriate balance between sample quality, data quality and impact on animals.

## 7. TAKE-HOME MESSAGES

1. In practice, most papers using the phrase “non-invasive DNA sampling” only comply to the medical definition of the term non-invasive, which is broader than the original definition proposed by Taberlet et al. [6] and is concerned only with the preservation of the physical integrity of the organism being sampled. We urge scientists using non-invasive DNA sampling methods to always state whether they refer to the definition by Taberlet et al. [6] *sensu stricto* or the medical definition of a non-invasive procedure (*sensu lato*).
2. We propose the new terms, “non-disruptive” and “minimally-disruptive” DNA sampling, to more appropriately address the potential behaviour, welfare and/or fitness effects of DNA sampling methods, as opposed to physical integrity (invasiveness in the medical sense). We can envisage situations in which the research aims are not impacted by the sampling approach to obtain DNA. However, researchers have an ethical obligation to minimise the impacts on the animals. Therefore, whenever possible, non-disruptive or minimally disruptive DNA sampling methods should be selected, in particular prior to experimental or observational studies measuring fitness or behaviour, as well as studies using techniques such as CMR where fitness or behaviour may affect results.
3. It may in some cases be better to use a physically invasive method (e.g. remote biopsy) that is minimally disruptive rather than a method that does not involve puncturing the skin but causes severe stress and has long-lasting effects (e.g. stressful capture for saliva swabbing).
4. More research is required to better understand the consequences of different live DNA sampling methods on behaviour, welfare and fitness in a variety of animal species and contexts.

## Data accessibility

The list of publications reviewed and the raw data used for the analyses are available in Supplementary Table 1.

## Supplementary material

The R codes and raw data are available online: 10.6084/m9.figshare.8397224

## Acknowledgements

The authors would like to thank Mehdi Mahjoob (http://mehdimahjoob.com), graphic designer, for his assistance with the preparation of Figure 3, the Environmental and Animal Sciences Writing Group at Unitec Institute of Technology, and particularly Rebecca Ladyman, for their valuable feedback on an earlier version of the manuscript. This review was initiated at Lincoln University (New Zealand) and we are grateful for the logistical support provided by this institution. Version 4 of this preprint has been reviewed and recommended by Peer Community In Ecology (https://doi.org/10.24072/pci.ecology.100029)

## Authors contribution

Conceptualised the idea: MCL, SB, RHC; recruited co-authors and organised literature review and writing workshops: MCL, SB; conducted the systematic review: MCL, SB; prepared the figures: SB; drafted and revised the manuscript MCL, RHC, NA, AB, KD, AK, JR, VRS, RS, WB, SB.

The overall author percentage contributions are as follow: MCL^25^, RHC^10^, KD^8^, NA^5^, AB^5^, AK^5^, JR^5^, VRS^5^, RS^5^, BW^2^, SB^25^.

## Conflict of interest disclosure

The authors of this preprint declare that they have no financial conflict of interest with the content of this article.

## Notes

#### Summary of Updates

This version has been reviewed and recommended by Peer Community In Ecology.

https://doi.org/10.6084/m9.figshare.8397224.v1

## References

1. Suarez RK, Moyes CD (2012) Metabolism in the age of “omes”. J Exp Biol 215: 2351–2357. Available: http://www.ncbi.nlm.nih.gov/pubmed/22723473. Accessed 1 December 2014.

2. Sboner A, Mu X, Greenbaum D (2011) The real cost of sequencing: higher than you think! Genome Biol 12: 125. Available: http://www.biomedcentral.com/content/pdf/gb-2011-12-8-125.pdf. Accessed 19 January 2015.

3. Hamilton MJ, Sadowsky MJ (2010) DNA Profiling in Ecology. Encyclopedia of Life Sciences. Major Reference Works. Chichester, UK: John Wiley & Sons, Ltd. Available: https://doi.org/10.1002/9780470015902.a0005454.pub2.

4. Hollands C (1986) The animals (scientific procedures) Act 1986. Lancet 328: 32–33. Available: http://linkinghub.elsevier.com/retrieve/pii/S0140673686925717.

5. Taberlet P, Luikart G (1999) Non-invasive genetic sampling and individual identification. Biol J Linn Soc 68: 41–55. Available: http://linkinghub.elsevier.com/retrieve/pii/S0024406699903292.

6. Taberlet P, Waits LP, Luikart G (1999) Noninvasive genetic sampling: look before you leap. Trends Ecol Evol 14: 323–327. Available: http://linkinghub.elsevier.com/retrieve/pii/S0169534799016377.

7. R Core Team (2018) R: A language and environment for statistical computing. R Foundation for Statistical Computing. Vienna, Austria. doi:10.1108/eb003648.

8. RStudio (2017) RStudio: Integrated development for R. [Online] RStudio, Inc, Boston, MA URL http://www.rstudio.com. doi:10.1007/978-81-322-2340-5.

9. Roy J, Vigilant L, Gray M, Wright E, Kato R, et al. (2014) Challenges in the use of genetic mark-recapture to estimate the population size of Bwindi mountain gorillas (Gorilla beringei beringei). Biol Conserv 180: 249–261. Available: http://dx.doi.org/10.1016/j.biocon.2014.10.011.

10. Sugimoto T, Aramilev V V., Kerley LL, Nagata J, Miquelle DG, et al. (2014) Noninvasive genetic analyses for estimating population size and genetic diversity of the remaining Far Eastern leopard (Panthera pardus orientalis) population. Conserv Genet 15: 521–532. Available: http://link.springer.com/10.1007/s10592-013-0558-8.

11. Garshelis DL (2006) On the allure of noninvasive genetic sampling-putting a face to the name. Ursus 17: 109–123. doi:10.2192/1537-6176(2006)17[109:OTAONG]2.0.CO;2.

12. Waldner T, Traugott M (2012) DNA-based analysis of regurgitates: a noninvasive approach to examine the diet of invertebrate consumers. Mol Ecol Resour 12: 669–675. Available: http://www.ncbi.nlm.nih.gov/pubmed/22443278. Accessed 23 January 2015.

13. Baumgardt JA, Goldberg CS, Reese KP, Connelly JW, Musil DD, et al. (2013) A method for estimating population sex ratio for sage-grouse using noninvasive genetic samples. Mol Ecol Resour 13: 393–402. Available: http://www.ncbi.nlm.nih.gov/pubmed/23347565. Accessed 14 August 2014.

14. Williams DA, Leach C, Hale AM, Karsten KB, Mujica E, et al. (2012) Development of tetranucleotide microsatellite loci and a non-invasive DNA sampling method for Texas horned lizards (Phrynosoma cornutum). Conserv Genet Resour 4: 43– 45. doi:10.1007/s12686-011-9469-5.

15. Murphy D, Noon B (1991) Coping with uncertainty in wildlife biology. J Wildl Manage 55: 773–782. Available: http://www.jstor.org/stable/3809531. Accessed 19 January 2015.

16. Hodges KE (2008) Defining the problem: terminology and progress in ecology. Front Ecol Environ 6: 35–42. Available: http://www.esajournals.org/doi/abs/10.1890/060108. Accessed 30 December 2014.

17. Herrando-Perez S, Brook BW, Bradshaw CJ a. (2014) Ecology needs a convention of nomenclature. Bioscience 64: 311–321. Available: http://bioscience.oxfordjournals.org/cgi/doi/10.1093/biosci/biu013. Accessed 19 January 2015.

18. Rorat A, Kachamakova-Trojanowska N, Jozkowicz A, Kruk J, Cocquerelle C, et al. (2014) Coelomocyte-Derived Fluorescence and DNA Markers of Composting Earthworm Species. J Exp Zool PART A-ECOLOGICAL Genet Physiol 321: 28–40. doi:10.1002/jez.1834.

19. Quynh NH, Inn KY, Amaël B, Yikweon J (2017) Efficient isolation method for high-quality genomic DNA from cicada exuviae. Ecol Evol 7: 8161–8169. Available: https://doi.org/10.1002/ece3.3398.

20. Richter A, Weinhold D, Robertson G, Young M, Edwards T, et al. (2013) More than an empty case: a non invasive technique for monitoring the Australian critically endangered golden sun moth, Synemon plana (Lepidoptera: Castniidae). J Insect Conserv 17: 529–536. doi:10.1007/s10841-012-9537-5.

21. Lefort M-C, Wratten S, Cusumano A, Varennes Y-D, Boyer S (2017) Disentangling higher trophic level interactions in the cabbage aphid food web using high-throughput DNA sequencing. Metabarcoding and Metagenomics 1: e13709. Available: https://mbmg.pensoft.net/articles.php?id=13709. Accessed 18 October 2017.

22. Madden AA, Barberán A, Bertone MA, Menninger HL, Dunn RR, et al. (2016) The diversity of arthropods in homes across the United States as determined by environmental DNA analyses. Mol Ecol 25: 6214–6224. Available: http://doi.wiley.com/10.1111/mec.13900.

23. Bienert F, De Danieli S, Miquel C, Coissac E, Poillot C, et al. (2012) Tracking earthworm communities from soil DNA. Mol Ecol 21: 2017–2030. Available: http://www.ncbi.nlm.nih.gov/pubmed/22250728. Accessed 9 November 2012.

24. Mächler E, Deiner K, Steinmann P, Altermatt F (2014) Utility of environmental DNA for monitoring rare and indicator macroinvertebrate species. Freshw Sci 33: 1174–1183. Available: http://www.bioone.org/doi/abs/10.1086/678128.

25. Baillon L, Pierron F, Oses J, Pannetier P, Normandeau E, et al. (2016) Detecting the exposure to Cd and PCBs by means of a non-invasive transcriptomic approach in laboratory and wild contaminated European eels (Anguilla anguilla). Environ Sci Pollut Res 23: 5431–5441. doi:10.1007/s11356-015-5754-2.

26. Janse M, Kappe AL, Van Kuijk BLM (2013) Paternity testing using the poisonous sting in captive white-spotted eagle rays Aetobatus narinari: a non-invasive tool for captive sustainability programmes. J Fish Biol 82: 1082–1085. doi:10.1111/jfb.12038.

27. Roques JAC, Abbink W, Geurds F, van de Vis H, Flik G (2010) Tailfin clipping, a painful procedure: Studies on Nile tilapia and common carp. Physiol Behav 101: 533–540. Available: http://linkinghub.elsevier.com/retrieve/pii/S0031938410002866.

28. Roffler GH, Talbot SL, Luikart G, Sage GK, Pilgrim KL, et al. (2014) Lack of sex-biased dispersal promotes fine-scale genetic structure in alpine ungulates. Conserv Genet 15: 837–851. doi:10.1007/s10592-014-0583-2.

29. Ditmer MA, Vincent JB, Werden LK, Tanner JC, Laske TG, et al. (2015) Bears show a physiological but limited behavioral response to unmanned aerial vehicles. Curr Biol 25: 2278–2283. Available: http://linkinghub.elsevier.com/retrieve/pii/S0960982215008271.

30. Woodruff SP, Johnson TR, Waits LP (2016) Examining the use of fecal pellet morphometry to differentiate age classes in Sonoran pronghorn. Wildlife Biol 22: 217–227. doi:10.2981/wlb.00209.

31. Brown VA, Willcox E V., Fagan KE, Bernard RF (2017) Identification of southeastern bat species using noninvasive genetic sampling of individual guano pellets. J Fish Wildl Manag 8: 632–639. Available: http://www.fwspubs.org/doi/10.3996/012017-JFWM-007.

32. Jedlicka JA, Vo A-TE, Almeida RPP (2016) Molecular scatology and high-throughput sequencing reveal predominately herbivorous insects in the diets of adult and nestling Western Bluebirds (Sialia mexicana) in California vineyards. Auk 134: 116–127. doi:10.1642/auk-16-103.1.

33. Llaneza L, García EJ, López-Bao JV (2014) Intensity of territorial marking predicts wolf reproduction: implications for wolf monitoring. PLoS One 9: e93015. Available: http://dx.plos.org/10.1371/journal.pone.0093015.

34. Brzeziński M, Romanowski J (2006) Experiments on sprainting activity of otters (Lutra lutra) in the Bieszczady Mountains, southeastern Poland / Observations des épreintes de la loutre (Lutra lutra) sur les montagnes du Bieszczady au sud-est de la Pologne. Mammalia 70: 58–63. Available: http://www.degruyter.com/view/j/mamm.2006.70.issue-1_2/mamm.2006.019/mamm.2006.019.xml. Accessed 5 January 2015.

35. Bonesi L, Hale M, Macdonald DW (2013) Lessons from the use of non-invasive genetic sampling as a way to estimate Eurasian otter population size and sex ratio. Acta Theriol (Warsz) 58: 157–168. doi:10.1007/s13364-012-0118-5.

36. Mannise N, Trovati RG, Duarte JMB, Maldonado JE, González S (2018) Using non–Invasive genetic techniques to assist in maned wolf conservation in a remnant fragment of the Brazilian Cerrado. Anim Biodivers Conserv 41: 315– 319.

37. Klütsch CFC, Thomas PJ (2018) Improved genotyping and sequencing success rates for North American river otter (Lontra canadensis). Eur J Wildl Res 64: 16. Available: http://link.springer.com/10.1007/s10344-018-1177-y.

38. Piaggio AJ, Cariappa CA, Straughan DJ, Neubaum MA, Dwire M, et al. (2016) A noninvasive method to detect Mexican wolves and estimate abundance. Wildl Soc Bull 40: 321–330. Available: http://doi.wiley.com/10.1002/wsb.659.

39. Stansbury CR, Ausband DE, Zager P, Mack CM, Miller CR, et al. (2014) A long-term population monitoring approach for a wide-ranging carnivore: Noninvasive genetic sampling of gray wolf rendezvous sites in Idaho, USA. J Wildl Manage 78: 1040–1049. Available: http://doi.wiley.com/10.1002/jwmg.736.

40. Wultsch C, Waits LP, Hallerman EM, Kelly MJ (2015) Optimizing collection methods for noninvasive genetic sampling of neotropical felids. Wildl Soc Bull 39: 403–412. Available: http://doi.wiley.com/10.1002/wsb.540.

41. Tsaparis D, Karaiskou N, Mertzanis Y, Triantafyllidis A (2015) Non-invasive genetic study and population monitoring of the brown bear (Ursus arctos) (Mammalia: Ursidae) in Kastoria region-Greece. J Nat Hist 49: 393–410. doi:10.1080/00222933.2013.877992.

42. Stansbury CR, Ausband DE, Zager P, Mack CM, Waits LP (2016) Identifying gray wolf packs and dispersers using noninvasive genetic samples. J Wildl Manage 80: 1408–1419. Available: http://doi.wiley.com/10.1002/jwmg.21136.

43. Karamanlidis AA, Stojanov A, de Gabriel Hernando M, Ivanov G, Kocijan I, et al. (2014) Distribution and genetic status of brown bears in FYR Macedonia: implications for conservation. Acta Theriol (Warsz) 59: 119–128. Available: http://link.springer.com/10.1007/s13364-013-0147-8.

44. Quinn TP, Wirsing AJ, Smith B, Cunningham CJ, Ching J (2014) Complementary use of motion-activated cameras and unbaited wire snares for DNA sampling reveals diel and seasonal activity patterns of brown bears (Ursus arctos) foraging on adult sockeye salmon (Oncorhynchus nerka). Can J Zool 92: 893– 903. Available: https://doi.org/10.1139/cjz-2014-0114.

45. Godwin BL, Albeke SE, Bergman HL, Walters A, Ben-David M (2015) Density of river otters (Lontra canadensis) in relation to energy development in the Green River Basin, Wyoming. Sci Total Environ 532: 780–790. Available: http://linkinghub.elsevier.com/retrieve/pii/S0048969715302552.

46. Cohen O, Barocas A, Geffen E (2013) Conflicting management policies for the Arabian wolf Canis lupus arabs in the Negev Desert: is this justified? ORYX 47: 228–236. doi:10.1017/S0030605311001797.

47. Anile S, Arrabito C, Mazzamuto MV, Scornavacca D, Ragni B (2012) A non-invasive monitoring on European wildcat (Felis silvestris silvestris Schreber, 1777) in Sicily using hair trapping and camera trapping: does scented lure work? HYSTRIX-ITALIAN J Mammal 23: 44–49. doi:10.4404/hystrix-23.2-4657.

48. Steyer K, Simon O, Kraus RHS, Haase P, Nowak C (2013) Hair trapping with valerian-treated lure sticks as a tool for genetic wildcat monitoring in low-density habitats. Eur J Wildl Res 59: 39–46. doi:10.1007/s10344-012-0644-0.

49. Yannic G, Broquet T, Strøm H, Aebischer A, Dufresnes C, et al. (2016) Genetic and morphological sex identification methods reveal a male-biased sex ratio in the Ivory Gull Pagophila eburnea. J Ornithol 157: 861–873. Available: https://doi.org/10.1007/s10336-016-1328-4.

50. Cullingham CI, Thiessen CD, Derocher AE, Paquet PC, Miller JM, et al. (2016) Population structure and dispersal of wolves in the Canadian Rocky Mountains. J Mammal 97: 839–851. Available: https://academic.oup.com/jmammal/article-lookup/doi/10.1093/jmammal/gyw015.

51. Dai Y, Lin Q, Fang W, Zhou X, Chen X (2015) Noninvasive and nondestructive sampling for avian microsatellite genotyping: a case study on the vulnerable Chinese Egret (Egretta eulophotes). Avian Res 6: 24. Available: http://www.avianres.com/content/6/1/24.

52. Barbosa S, Pauperio J, Searle JB, Alves PC (2013) Genetic identification of Iberian rodent species using both mitochondrial and nuclear loci: application to noninvasive sampling. Mol Ecol Resour 13: 43–56. Available: http://doi.wiley.com/10.1111/1755-0998.12024.

53. Malekian M, Sadeghi P, Goudarzi F (2018) Assessment of environmental DNA for detection of an imperiled Amphibian, the luristan newt (Neurergus kaiseri, Schmidt 1952). Herpetol Conserv Biol 13: 175–182.

54. Ponjoan A, Bota G, De La Morena ELG, Morales MB, Wolff A, et al. (2008) Adverse effects of capture and handling little bustard. J Wildl Manage 72: 315–319. Available: http://www.bioone.org/doi/abs/10.2193/2006-443.

55. Harcourt RG, Turner E, Hall A, Waas JR, Hindell M (2010) Effects of capture stress on free-ranging, reproductively active male Weddell seals. J Comp Physiol A 196: 147–154. Available: http://link.springer.com/10.1007/s00359-009-0501-0.

56. Seguel M, Paredes E, Pavés H, Gottdenker NL (2014) Capture-induced stress cardiomyopathy in South American fur seal pups (Arctophoca australis gracilis). Mar Mammal Sci 30: 1149–1157. Available: http://doi.wiley.com/10.1111/mms.12079.

57. Hollenbeck N, Scheel D, Gravley MC, Sage GK, Toussaint R, et al. (2017) Use of Swabs for Sampling Epithelial Cells for Molecular Genetics Analyses in Enteroctopus. Am Malacol Bull 35: 145–157.

58. Fiorito G, Affuso A, Basil J, Cole A, de Girolamo P, et al. (2015) Guidelines for the care and welfare of cephalopods in research – A consensus based on an initiative by cephRes, FELASA and the Boyd Group. Lab Anim 49: 1–90. Available: http://journals.sagepub.com/doi/10.1177/0023677215580006.

59. Li J, Cui Y, Jiang J, Yu J, Niu L, et al. (2017) Applying DNA barcoding to conservation practice: a case study of endangered birds and large mammals in China. Biodivers Conserv 26: 653–668. doi:10.1007/s10531-016-1263-y.

60. Porco D, Rougerie R, Deharveng L, Hebert P (2010) Coupling non-destructive DNA extraction and voucher retrieval for small soft-bodied Arthropods in a high-throughput context: the example of Collembola. Mol Ecol Resour 10: 942– 945. Available: http://doi.wiley.com/10.1111/j.1755-0998.2010.2839.x.

61. Wisely SM, Maldonado JE, Fleische RC (2004) A technique for sampling ancient DNA that minimizes damage to museum specimens. Conserv Genet 5: 105–107. Available: http://link.springer.com/10.1023/B:COGE.0000014061.04963.da.

62. Koczur LM, Williford D, DeYoung RW, Ballard BM (2017) Bringing back the dead: Genetic data from avian carcasses. Wildl Soc Bull 41: 796–803. Available: http://doi.wiley.com/10.1002/wsb.823.

63. Monteiro NM, Silva RM, Cunha M, Antunes A, Jones AG, et al. (2014) Validating the use of colouration patterns for individual recognition in the worm pipefish using a novel set of microsatellite markers. Mol Ecol Resour 14: 150–156. doi:10.1111/1755-0998.12151.

64. Ringler E (2018) Testing skin swabbing for DNA sampling in dendrobatid frogs. Amphibia-Reptilia 39: 245–251. Available: http://booksandjournals.brillonline.com/content/journals/10.1163/15685381-17000206.

65. Player D, Lausen C, Zaitlin B, Harrison J, Paetkau D, et al. (2017) An alternative minimally invasive technique for genetic sampling of bats: Wing swabs yield species identification. Wildl Soc Bull 41: 590–596. Available: http://doi.wiley.com/10.1002/wsb.803.

66. Jones SP, Kennedy SW (2015) Feathers as a source of RNA for genomic studies in avian species. Ecotoxicology 24: 55–60. Available: http://link.springer.com/10.1007/s10646-014-1354-z.

67. Ford B, Govindarajulu P, Larsen K, Russello M (2017) Evaluating the efficacy of non-invasive genetic sampling of the Northern Pacific rattlesnake with implications for other venomous squamates. Conserv Genet Resour 9: 13–15. doi:10.1007/s12686-016-0606-z.

68. Barbosa S, Paupério J, Herman JS, Ferreira CM, Pita R, et al. (2017) Endemic species may have complex histories: within-refugium phylogeography of an endangered Iberian vole. Mol Ecol 26: 951–967. Available: http://doi.wiley.com/10.1111/mec.13994.

69. du Toit Z, Grobler JP, Kotze A, Jansen R, Dalton DL (2017) Scale samples from Temminck’s ground pangolin (Smutsia temminckii): a non-invasive source of DNA. Conserv Genet Resour 9: 1–4. doi:10.1007/s12686-016-0602-3.

70. Miller-Keane, O’Toole MT (2005) Encyclopedia and dictionary of medicine, nursing, and allied health. Free Dict. doi:10.1097/00001610-199306000-00012.

71. Kauffman LK, Bjork JK, Gallup JM, Boggiatto PM, Bellaire BH, et al. (2014) Early detection of Brucella canis via quantitative polymerase chain reaction analysis. Zoonoses Public Health 61: 48–54. doi:10.1111/zph.12041.

72. Reinardy HC, Skippins E, Henry TB, Jha AN (2013) Assessment of DNA damage in sperm after repeated non-invasive sampling in zebrafish Danio rerio. J Fish Biol 82: 1074–1081. doi:10.1111/jfb.12042.

73. Michael AM, Kirsten MP (2004) Clarifying the effect of toe clipping on frogs with Bayesian statistics. J Appl Ecol 41: 780–786. Available: http://ejournals.ebsco.com/direct.asp?ArticleID=409490A1B086D30990C8.

74. Marschalek D a., Jesu J a., Berres ME (2013) Impact of non-lethal genetic sampling on the survival, longevity and behaviour of the Hermes copper (Lycaena hermes) butterfly. Insect Conserv Divers 6: 658–662. Available: http://doi.wiley.com/10.1111/icad.12024. Accessed 14 August 2014.

75. Oi CA, López-Uribe MM, Cervini M, Del Lama MA (2013) Non-lethal method of DNA sampling in euglossine bees supported by mark–recapture experiments and microsatellite genotyping. J Insect Conserv 17: 1071–1079. Available: http://link.springer.com/10.1007/s10841-013-9582-8. Accessed 14 August 2014.

76. Hamm CA, Aggarwal D, Landis DA (2010) Evaluating the impact of non-lethal DNA sampling on two butterflies, Vanessa cardui and Satyrodes eurydice. J Insect Conserv 14: 11–18. doi:10.1007/s10841-009-9219-0.

77. Vila M, Auger-Rozenberg M a., Goussard F, Lopez-Vaamonde C (2009) Effect of non-lethal sampling on life-history traits of the protected moth Graellsia isabelae (Lepidoptera: Saturniidae). Ecol Entomol 34: 356–362. Available: http://doi.wiley.com/10.1111/j.1365-2311.2008.01084.x. Accessed 14 August 2014.

78. Doerr L, Richardson K, Ewen J, Armstrong D (2017) Effect of supplementary feeding on reproductive success of hihi (stitchbird, Notiomystis cincta) at a mature forest reintroduction site. N Z J Ecol 41: 34–40. Available: http://newzealandecology.org/nzje/3295.

79. Brekke P, Benner PM, Santure AW, Ewen JG (2011) High genetic diversity in the remnant island population of hihi and the genetic consequences of re-introduction. Mol Ecol 20: 29–45. Available: http://doi.wiley.com/10.1111/j.1365-294X.2010.04923.x.

80. Bearzi G (2000) First report of a common dolphin (Delphinus delphis) death following penetration of a biopsy dart. J Cetacean Res Manag 2: 217–221. Available: http://onlinelibrary.wiley.com/doi/10.1111/j.1748-7692.1996.tb00302.x/full. Accessed 19 January 2015.

81. Gemmell N, Majluf P (1997) Projectile biopsy sampling of fur seals. Mar Mammal Sci 13: 512–516. Available: http://www.researchgate.net/publication/249473950_PROJECTILE_BIOPSY_SAMPLING_OF_FUR_SEALS/file/60b7d52273289d686b.pdf. Accessed 19 January 2015.

82. Krützen M, Barré L (2002) A biopsy system for small cetaceans: darting success and wound healing in Tursiops spp. Mar Mammal Sci 18: 863–878. Available: http://onlinelibrary.wiley.com/doi/10.1111/j.1748-7692.2002.tb01078.x/abstract. Accessed 19 January 2015.

83. Brown MW, Kraus SD, D.E. G (1991) Reaction of North Atlantic right whales (Eubalaena glacialis) to skin biopsy sampling for genetic and pollutant analysis. Report of the International Whaling Commission Special Issue. pp. 1381–1389. doi:i0022-541X-69-3-1171-Brown2.

84. Barrett-Lennard L (1996) A cetacean biopsy system using lightweight pneumatic darts, and its effect on the behavior of killer whales. Mar Mammal Sci 12: 14–27. Available: http://onlinelibrary.wiley.com/doi/10.1111/j.1748-7692.1996.tb00302.x/full. Accessed 19 January 2015.

85. Gauthier J, Sears R (1999) Behavioral response of four species of balaenopterid whales to biopsy sampling. Mar Mammal Sci 15: 85–101. Available: http://onlinelibrary.wiley.com/doi/10.1111/j.1748-7692.1999.tb00783.x/abstract. Accessed 19 January 2015.

86. Parris KM, McCall SC, McCarthy MA, Minteer BA, Steele K, et al. (2010) Assessing ethical trade-offs in ecological field studies. J Appl Ecol 47: 227–234. doi:10.1111/j.1365-2664.2009.01755.x.

87. Magoun AJ, Long CD, Schwartz MK, Pilgrim KL, Lowell RE, et al. (2011) Integrating motion-detection cameras and hair snags for wolverine identification. J Wildl Manage 75: 731–739. Available: http://doi.wiley.com/10.1002/jwmg.107.

88. Rovang S, Nielsen SE, Stenhouse G (2015) In the trap: detectability of fixed hair trap DNA methods in grizzly bear population monitoring. Wildlife Biol 21: 68–79. Available: http://www.bioone.org/doi/10.2981/wlb.00033.

89. Henry P, Henry A, Russello MA (2011) A noninvasive hair sampling technique to obtain high quality DNA from elusive small mammals. J Vis Exp. Available: http://www.jove.com/index/Details.stp?ID=2791.

90. Banks SC, Hoyle SD, Horsup A, Sunnucks P, Taylor AC (2003) Demographic monitoring of an entire species (the northern hairy-nosed wombat, Lasiorhinus krefftii) by genetic analysis of non-invasively collected material. Anim Conserv 6: 101–107. Available: http://doi.wiley.com/10.1017/S1367943003003135.

91. Dalén L, Götherström A (2007) Recovery of DNA from footprints in the snow. Can Field-Naturalist 121: 321–324. Available: http://canadianfieldnaturalist.ca/index.php/cfn/article/viewArticle/482. Accessed 19 January 2015.

92. Nichols R V, Königsson H, Danell K, Spong G (2012) Browsed twig environmental DNA: diagnostic PCR to identify ungulate species. Mol Ecol Resour 12: 983–989. Available: http://www.ncbi.nlm.nih.gov/pubmed/22813481. Accessed 29 January 2014.

93. Prunier J, Kaufmann B, Grolet O, Picard D, Pompanon F, et al. (2012) Skin swabbing as a new efficient DNA sampling technique in amphibians, and 14 new microsatellite markers in the alpine newt (Ichthyosaura alpestris). Mol Ecol Resour 12: 524–531. Available: http://doi.wiley.com/10.1111/j.1755-0998.2012.03116.x.

94. Morinha F, Travassos P, Carvalho D, Magalhaes P, Cabral JA, et al. (2014) DNA sampling from body swabs of terrestrial slugs (Gastropoda: Pulmonata): a simple and non-invasive method for molecular genetics approaches. J Molluscan Stud 80: 99–101. Available: https://academic.oup.com/mollus/article-lookup/doi/10.1093/mollus/eyt045.

95. Uno R, Kondo M, Yuasa T, Yamauchi K, Tsuruga H, et al. (2012) Assessment of genotyping accuracy in a non-invasive DNA-based population survey of Asiatic black bears (Ursus thibetanus): lessons from a large-scale pilot study in Iwate prefecture, northern Japan. Popul Ecol 54: 509–519. Available: http://link.springer.com/10.1007/s10144-012-0328-3. Accessed 14 August 2014.

96. Collins R a., Armstrong KF, Holyoake AJ, Keeling S (2012) Something in the water: biosecurity monitoring of ornamental fish imports using environmental DNA. Biol Invasions 15: 1209–1215. Available: http://link.springer.com/10.1007/s10530-012-0376-9. Accessed 7 March 2013.

97. Kowarski K (2014) Effects of remote biopsy sampling on long-finned pilot whales (Globicephala melas) in Nova Scotia. Aquat Mamm 40: 117–125. Available: http://aquaticmammalsjournal.org/index.php?option=com_content&view=article&id=680:effects-of-remote-biopsy-sampling-on-long-finned-pilot-whales-globicephala-melas-in-nova-scotia&catid=55&Itemid=157.

98. Pagano AM, Peacock E, McKinney MA (2014) Remote biopsy darting and marking of polar bears. Mar Mammal Sci 30: 169–183. Available: http://doi.wiley.com/10.1111/mms.12029.

99. Boyer S, Wratten SD, Holyoake A, Abdelkrim J, Cruickshank RH (2013) Using next-generation sequencing to analyse the diet of a highly endangered land snail (Powelliphanta augusta) feeding on endemic earthworms. PLoS One 8: e75962. Available: http://dx.plos.org/10.1371/journal.pone.0075962. Accessed 29 September 2013.

100. Lefort M-C, Boyer S, De Romans S, Glare T, Armstrong K, et al. (2014) Invasion success of a scarab beetle within its native range: host range expansion versus host-shift. PeerJ 2: e262. Available: https://peerj.com/articles/262. Accessed 26 February 2014.

101. Waterhouse BR, Boyer S, Wratten SD (2014) Pyrosequencing of prey DNA in faeces of carnivorous land snails to facilitate ecological restoration and relocation programmes. Oecologia 175: 737–746. Available: http://link.springer.com/10.1007/s00442-014-2933-7. Accessed 29 March 2014.

102. Bateson M (2016) Cumulative stress in research animals: Telomere attrition as a biomarker in a welfare context? BioEssays 38: 201–212. doi:10.1002/bies.201500127.

103. Vucicevic M, Stevanov-Pavlovic M, Stevanovic J, Bosnjak J, Gajic B, et al. (2013) Sex determination in 58 bird species and evaluation of CHD gene as a universal molecular marker in bird sexing. Zoo Biol 32: 269–276. Available: http://www.ncbi.nlm.nih.gov/pubmed/22553188. Accessed 19 January 2015.

104. Hebert PDN, Penton EH, Burns JM, Janzen DH, Hallwachs W (2004) Ten species in one: DNA barcoding reveals cryptic species in the neotropical skipper butterfly Astraptes fulgerator. Proc Natl Acad Sci U S A 101: 14812–14817. Available: http://www.pubmedcentral.nih.gov/articlerender.fcgi?artid=522015&tool=pmcentrez&rendertype=abstract.

105. Fumanal B, Martin J-F, Bon MC (2005) High through-put characterization of insect morphocryptic entities by a non-invasive method using direct-PCR of fecal DNA. J Biotechnol 119: 15–19. Available: http://www.ncbi.nlm.nih.gov/pubmed/15961176. Accessed 14 August 2014.

106. Creel S (2001) Social dominance and stress hormones. Trends Ecol Evol 16: 491–497. Available: http://linkinghub.elsevier.com/retrieve/pii/S0169534701022273.

107. Suleman M, Wango E (2004) Physiologic manifestations of stress from capture and restraint of free-ranging male African green monkeys (Cercopithecus aethiops). J Zoo Wildl Med 35: 20–24. Available: http://www.bioone.org/doi/abs/10.1638/01-025. Accessed 19 January 2015.

108. Thomson J a., Heithaus MR (2014) Animal-borne video reveals seasonal activity patterns of green sea turtles and the importance of accounting for capture stress in short-term biologging. J Exp Mar Bio Ecol 450: 15–20. Available: http://linkinghub.elsevier.com/retrieve/pii/S0022098113003596. Accessed 1 December 2014.

109. Robinson S, Waits L, Martin I (2009) Estimating abundance of American black bears using DNA-based capture-mark-recapture models. Ursus 20: 1–11. Available: http://www.bioone.org/doi/abs/10.2192/08GR022R.1. Accessed 19 January 2015.

110. Nelson G, Graves B (2004) Anuran population monitoring: comparison of the north American Amphibian monitoring program’s calling index with mark-recapture estimates for Rana clamitans. J Herpetol 38: 355–359. Available: http://www.bioone.org/doi/abs/10.1670/22-04A. Accessed 19 January 2015.

111. Boulanger J, Stenhouse G, Munro R (2004) Sources of heterogeneity bias when DNA mark-recapture sampling methods are applied to grizzly bear (ursus arctos) populations. J Mammal 85: 618–624. Available: https://academic.oup.com/jmammal/article-lookup/doi/10.1644/BRB-134.

112. Lampa S, Henle K, Klenke R, Hoehn M, Gruber B (2013) How to overcome genotyping errors in non-invasive genetic mark-recapture population size estimation-A review of available methods illustrated by a case study. J Wildl Manage 77: 1490–1511. Available: http://doi.wiley.com/10.1002/jwmg.604. Accessed 11 August 2014.

113. Modave E, MacDonald AJ, Sarre SD (2017) A single mini-barcode test to screen for Australian mammalian predators from environmental samples. Gigascience 6: 1–13. doi:10.1093/gigascience/gix052.

114. Duenas JF, Cruickshank R, Ross J (2015) Optimisation of a microsatellite panel for the individual identification of brushtail possums using low template DNA. N Z J Ecol 39: 93–102.

115. Huang H, Wang H, Li L, Wu Z, Chen J (2014) Genetic diversity and population demography of the Chinese crocodile lizard (Shinisaurus crocodilurus) in China. PLoS One 9. doi:10.1371/journal.pone.0091570.

116. Proffitt KM, Goldberg JF, Hebblewhite M, Russell R, Jimenez BS, et al. (2015) Integrating resource selection into spatial capture-recapture models for large carnivores. ECOSPHERE 6. doi:10.1890/ES15-00001.1.

117. López H, Contreras-Díaz HG, Oromí P, Juan C (2007) Delimiting species boundaries for endangered Canary Island grasshoppers based on DNA sequence data. Conserv Genet 8: 587–598. doi:10.1007/s10592-006-9199-5.

